# Genetic Diversity of *Candidatus* Liberibacter asiaticus Based on Four Hypervariable Genomic Regions in China

**DOI:** 10.1101/2022.03.21.485244

**Authors:** Fanglan Gao, Bo Wu, Chengwu Zou, Yixue Bao, Dean Li, Wei Yao, Charles A. Powell, Muqing Zhang

**Affiliations:** State Key Laboratory for Conservation and Utilization of Sub-tropical Bio-Agricultural Resources, Guangxi University, Guangxi, 530005, China; School of Computing, Clemson University, 821 McMillan Rd, Clemson, SC 29631, USA; Fruit Experimental Station, Agricultural and Rural Department of Guangxi, Nanning, 530002, China; IRREC, IFAS, University of Florida, Fort Pierce, FL 34945, USA

**Keywords:** *Candidatus* Liberibacter asiaticus, Huanglongbing (HLB), Hypervariable area, SNP, Genetic diversity

## Abstract

Huanglongbing (HLB; greening disease), caused by *Candidatus* Liberibacter asiaticus (*C*Las), is the most damaging citrus disease worldwide. The disease has spread throughout the citrus-producing regions of Guangxi, Guangdong, Fujian, and others in China. A total of 1,789 HLB-like symptomatic or asymptomatic samples were collected from the Guangxi and Fujian provinces of China to decipher the genetic diversity of *C*Las and its correlation with pathogenicity and host range. The disease was the most severe in orange and the least in pomelo. *C*Las bacteria associated with the specific geographical and citrus variety infected more than 50% of the HLB-like symptomatic samples. We identified a total of 6,286 minor variations by comparing 35 published *C*Las genomes and observed a highly heterogeneous variation distribution across the genome. Highly diverse regions, including two prophages, were generally more unstable and prone to lose. In the collected *C*Las strains, four highly diverse non-prophage segments and three prophage segments in the prophage region were chosen for PCR amplification and genotyping. Four hypervariable regions were used to decipher *C*Las diversity. A total of 100 strains were divided into four groups, of which 90 strains were clustered into two clades with 15 reported reference genomes, while 10 were grouped in two clades separately from the reported genomes. This study provides insight into the molecular characteristics and genetic variations of different *C*Las strains in China and might help develop HLB prevention and control strategies.

**Importance:** The objective of the present study is to characterize the hypervariable genomic region sequence, decipher the genetic diversity of *C*Las strains from China, understand how various strains differ, and lay the foundation for developing HLB prevention and control strategies.

## Introduction

Citrus huanglongbing (HLB), an invasive and severe disease, is devastating to most citrus species worldwide (1). The fruit drop and poor quality caused by HLB epidemics has severely harmed the commercial value of citrus cultivars in China since the disease was discovered in the late 1800s (2). HLB is associated with three gram-negative, phloem-residing bacteria in the genus *Candidatus* Liberibacter (3), which are vectored by Asian citrus psyllids (4, 5). *Candidatus* Liberibacter asiaticus (*C*Las) infects all planted citrus varieties. Severely HLB-infected citrus trees show stunted yellow shoots, blotchy mottle on the leaves, sparse foliation, and even die back, losing their economic vitality (6, 7). The symptoms in the field are diverse and highly detrimental to HLB detection, prevention, and control. The most effective management of citrus HLB includes removing pathogens by eradicating infected trees, cultivating pathogen-free nurseries, and pesticide control of Asian citrus psyllid (8). Almost all citrus varieties are susceptible to HLB, and resistance against this disease has not been developed. Genetic variation in conserved genes cannot distinguish closely related isolates. Therefore, molecular marker methods might be combined to distinguish different isolates and identify the genetic diversity of *C*Las (9).

The genetic diversity of *C*Las isolates is deduced from different citrus varieties, geographical origins, population structure, and evolution. When *C*Las populations of different citrus species were compared, citrus genotypes drove pathogen genetic diversity. *C*Las strains are classified by analyzing the high-variation sequences of the conserved regions of pathogens and combining their infection characteristics and phenotypic symptoms to differentiate strains (10). The *C*Las pathogen is detected based on the relationship between bacteria and symptomatic performance at an early stage (3). In Florida, the HLB symptoms in citrus plants affected by HLB are associated with specific *C*Las populations. The dynamic change in the citrus endophytic bacterial population can also affect the *C*Las titer, closely related to the severity of symptoms (11). The metabolic components of plants also play a crucial role in the characteristics of pathogenic bacterial infection. By constructing genomic metabolic models of A4, FC17, GxPSY, IsI-1, Psy62, and Ycpsy, host interactions with specific strains are dependent on host metabolic phenotypes and primarily affected by nutrient deficiencies (L-proline, L-serine, and L-arginine) (12, 13).

More genomic loci have focused on *C*Las population variation (14, 15). This research aimed to characterize the new hypervariable regions in the reported genome for deciphering genomic diversity of *C*Las from Guangxi and Fujian, and to detect the new strains associated with citrus varieties, symptoms and geographical locations.

## Results

### Genetic Variations Among 35 *C*Las Genomes

To identify genomic segments with high diversity and universal presence in *C*Las isolates, we analyzed 35 published *C*Las genomes from nine countries, including 19 from the United States and 9 from China (Supplementary Table S1). Approximately 14.6% (185 kb) of the reference *C*Las strain GXPSY genome (1,268, 237bp) was missing in at least two genomes, including ~81.5kb in the two prophages and ~10.3kb in the non-prophage regions (Figure 1). The genomes had 6,285 small variations, including 6,012 single nucleotide variations (SNVs) and 273 small indels. The minor alleles of 2,160 of the variations were found in at least two genomes, while the remaining 4,125 were only supported by one. Both SNVs and indels had highly heterogeneous variations with significantly (p<0.001 by the two-tailed *t*-test) higher variation density in prophage regions (23.3 variations/kb) than in non-prophage regions (3.7 variations/kb). A high density of variations was found in three duplicate regions harboring rRNAs in the reference genome located at 398,493 bp ~ 403,387bp, 770,848 bp ~ 775,747bp, and 838,830bp ~ 843,728bp, most likely due to the assembly errors with next-generation sequencing data. In the protein-coding regions, a total of 3,779 variations were annotated, including 992 synonymous, 2,448 missense substitutions, and 117 frameshift indels (Supplementary Table S2). The prophage genes had significantly (*p*<0.001) more synonymous and missense variations than the non-prophage genes. *C*Las genotyping was performed on four non-prophage genomic segments with relatively high genetic diversity and universal presence in the 35 genomes (Figure 1), including CP004005.1: 1,096,498~1,097,081, CP004005.1: 661,737 ~ 662,418, CP004005.1: 464946 ~ 465629, and CP004005.1: 1174796 ~ 1175556.

**Figure 1:**
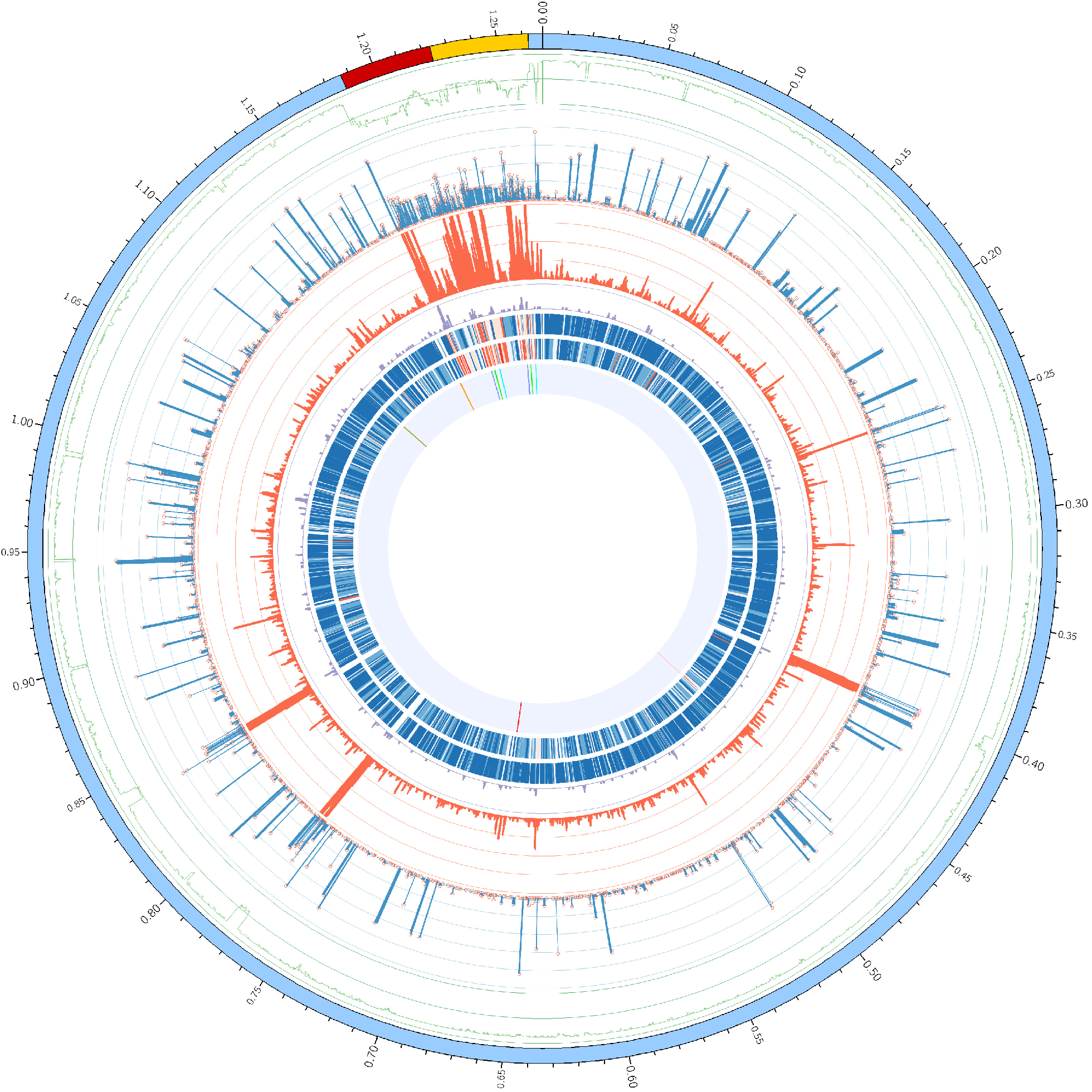
Circular graph depicting the distribution of variants in the 35 *C*Las genomes. From outer to inner: the ideogram of the reference GXPSY genome, with blue denoting non-prophage regions, red and orange regions denoting the two prophages, and the scale marks (Mb) indicating the coordinates on the reference; the line plot (green) depicts the number of *C*Las genomes in which the homologs of the local 1kb segments (overlapped by 500bp) were present; the distribution of the 6,285 variations (red hollow circles) across the reference and their observation times are represented by the height of the lines below the circles in the 35 *C*Las genomes; and the two histogram lines show the distributions of SNVs (red) and small indels (violet) in 1kb windows overlapped by 500bp across the reference, in which ≥50 and ≥10 values were shown as 50 and 10, correspondingly. The following two lines depict the densities of synonymous (outer lane) and missense (inner lane) SNVs from the lowest (dark blue) to the highest (dark red); the innermost ring indicates the locations of all the segments selected for PCR amplification and genotyping.

We performed the phylogenetic analysis using both neighbor-joining and Bayesian inference based on the 1,954 variations successfully genotyped in the 35 genomes. Nine *C*Las strains from China were classified into three distinct clades, at least two of which were introduced into the United States (Figure 2). The phylogenetic tree also confirmed that MEX8 (from Mexico) and COFLP (from Colombia) were related to strains in the United States. At the same time, 9PA (from Brazil) was obtained separately from China or other Asian areas.

**Figure 2:**
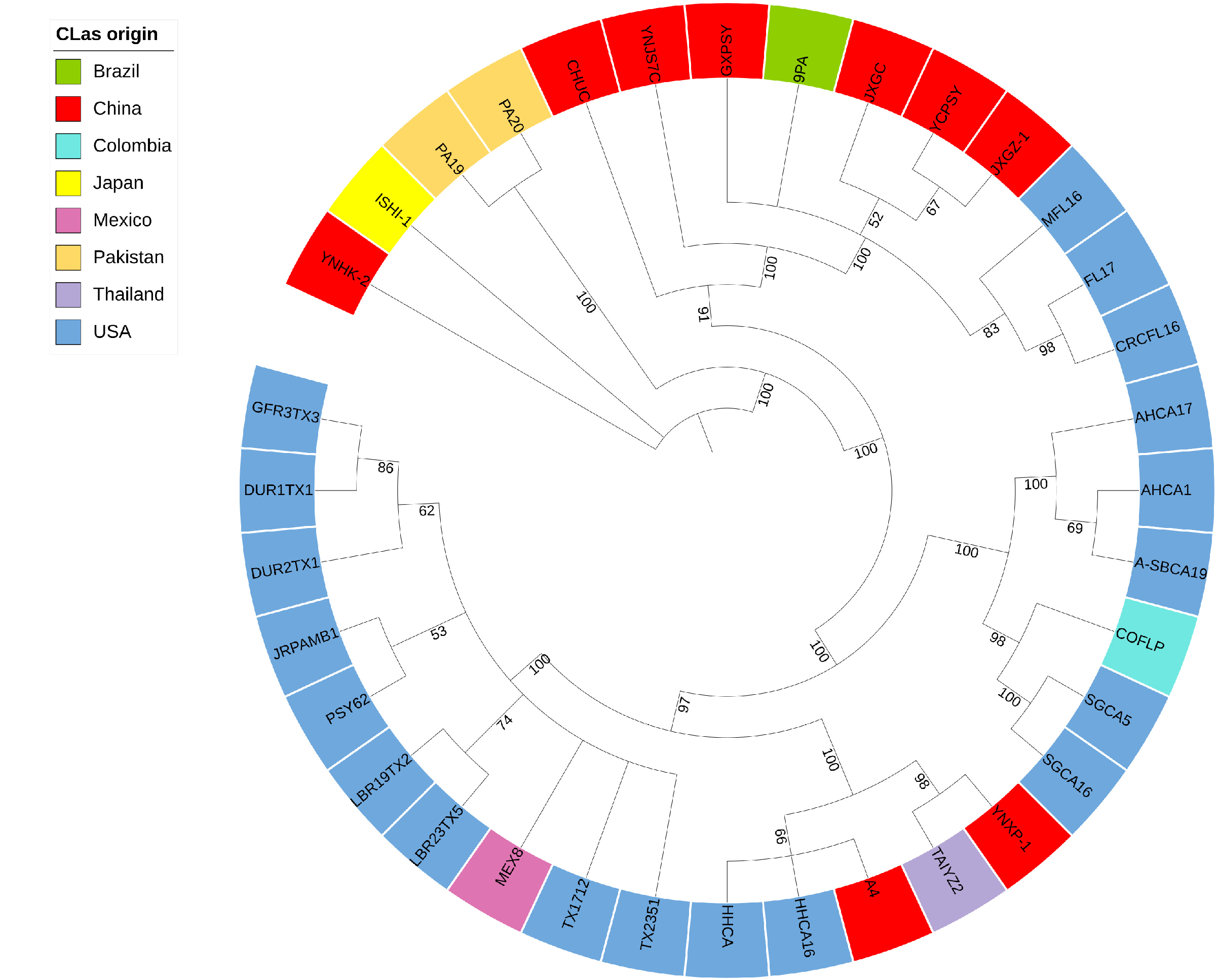
Phylogenetic tree based on 1,954 variations with available genotypes in all Las genomes. The phylogenetic tree was constructed using neighbor-joining (NJ) and Bayesian inference (BI). Only branches with ≥50% bootstrap support (NJ) and with ≥50% posterior probability (BI) are shown in the graph. The numbers on the branches represent the posterior probabilities (%) calculated using Bayesian inference.

### HLB Incidence in Citrus Groves in Guangxi and Fujian, China

A total of 1,789 samples were collected from 18 citrus groves in Fujian and Guangxi, with 1,365 (76.34%) being randomly collected and 423 (23.66%) being HLB-like symptomatic leaves. The highest percentage of *C*Las positive in randomly collected samples was 17% in Liuzhou, followed by 14.29% in Baise, 13.49% in Hezhou, and 13.21% in Wuzhou. However, the PI was over 20% in Wuzhou. HLB-like symptomatic leaves had a higher *C*Las positive percentage of 64.77% and a pathogen index (PI) of 45.32% than randomly collected samples (Table 1).

**Table 1.** Incidence of *C*Las in prominent citrus-producing areas in Guangxi and Fujian.

| Sampling<br>method | Sampling<br>sites | Tested<br>samples | PCR-positive<br>sample | PCR-positive<br>( % ) | CT value | PI (%) |
| --- | --- | --- | --- | --- | --- | --- |
| Randomly<br>sampling | Yulin, GX | 26 | 3 | 11.54 | 38.78±3.37 | 5.77 |
|  | Hezhou, GX | 126 | 17 | 13.49 | 39.28±3.27 | 4.30 |
|  | Laibin, GX | 65 | 5 | 7.69 | 39.17±3.24 | 4.62 |
|  | Guilin, GX | 408 | 29 | 7.11 | 39.51±2.32 | 2.68 |
|  | Wuzhou, GX | 106 | 14 | 13.21 | 36.26±5.34 | 20.83 |
|  | Liuzhou, GX | 100 | 17 | 17.00 | 38.69±2.99 | 6.25 |
|  | Hechi, GX | 100 | 6 | 6.00 | 39.53±2.04 | 2.50 |
|  | Baise, GX | 14 | 2 | 14.29 | 38.58±3.51 | 7.16 |
|  | Nanning, GX | 203 | 7 | 3.45 | 39.77±1.35 | 1.23 |
|  | Sanming, FJ | 217 | 15 | 6.91 | 39.13±3.40 | 4.44 |
|  | **Sub-total** | **1365** | **115** | **8.42** | **38.79±3.38** | **5.85** |
| HLB-like<br>sampling | Guigang, GX | 75 | 55 | 73.33 | 29.08±6.07 | 56.25 |
|  | Guilin, GX | 63 | 39 | 61.90 | 33.55±7.24 | 33.11 |
|  | Yulin, GX | 11 | 11 | 100.00 | 30.16±4.11 | 47.73 |
|  | Laibin, GX | 15 | 8 | 53.33 | 32.85±7.88 | 41.67 |
|  | Nanning, GX | 147 | 84 | 57.14 | 33.76±6.87 | 31.63 |
|  | Liuzhou, GX | 26 | 24 | 92.31 | 27.30±5.92 | 66.35 |
|  | Beihai, GX | 11 | 9 | 81.82 | 29.20±5.22 | 56.82 |
|  | Sanming, FJ | 75 | 44 | 58.67 | 33.00±6.63 | 36.67 |
|  | **Sub-total** | **423** | **274** | **64.77** | **31.16±5.83** | **45.32** |

A total of 423 HLB-like symptomatic samples were collected from four citrus varieties: mandarin (136), orange (64), pomelo (42), and tangerine (155). HLB affected all citrus varieties, ranging from 54.76% (pomelo) to 75.00% (orange). PI was highest in orange with the lowest CT value, while it was lowest in pomelo (Table 2). The typical HLB symptoms observed in the HLB-like symptomatic samples included yellowing, blotchy mottle, and Zn-deficient-like or asymptomatic leaves. Compared to yellowing and asymptomatic samples, blotchy mottle and Zn-deficient-like samples had a higher PCR-positive percentage and PI, but a lower CT value (Table 2).

**Table 2.**
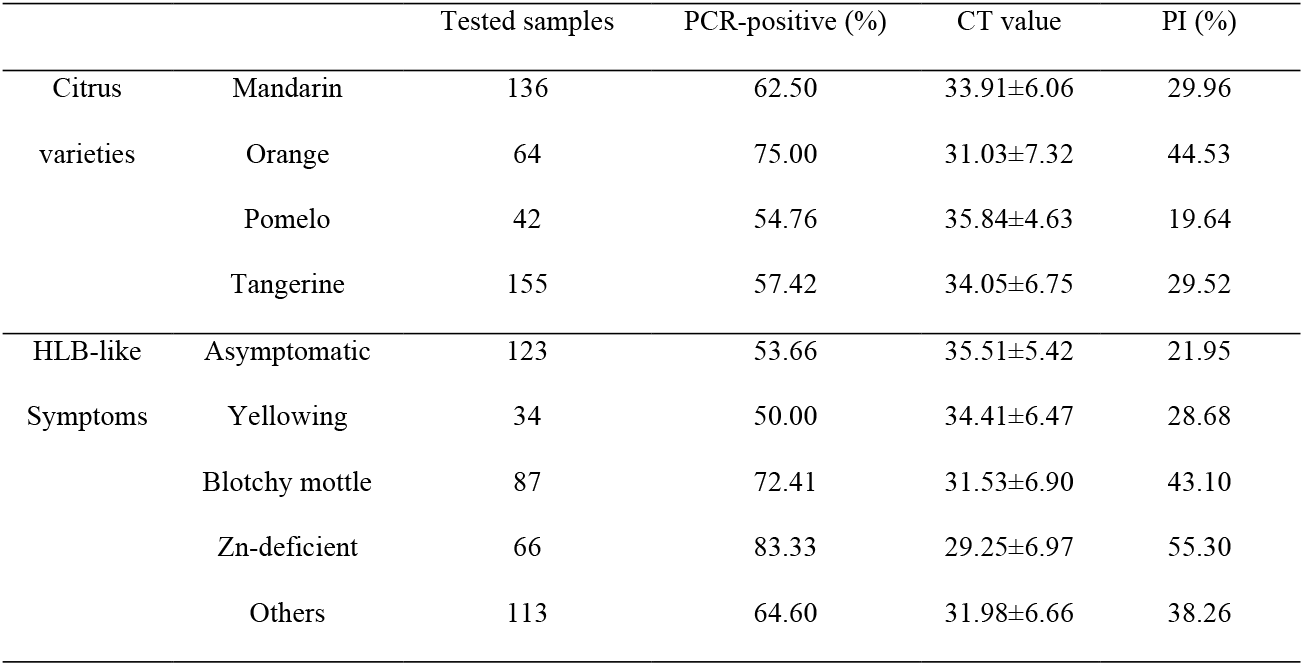
*C*Las variations in citrus varieties and HLB-like symptom types.

### *C*Las Population Dynamics in HLB-Infected Citrus Plants in China

Amplicons of the expected size were amplified and sequenced using the seven designed primers (Table 3, Supplementary S1). Only one individual base in a single-case mutation was detected in 69 samples from primer 124/805; nine in 57 from primer 121/804; and twenty-four in 62 from primer 230/1048 (Supplementary S3). The other four primers, including 465/1048, 141/901, 475/1055, and 309/1044, had high specificity and abundant mutation loci in the amplified regions, making them suitable for genotypic and phylogenetic analysis (Supplementary S2). Total 149 (43.44%) of the 343 *C*Las-positive DNA samples from China contained at least one of four hypervariable genomic regions (HGR), including 120 (80.54%) HGR-I from primer 465/1048, 126 (84.56%) HGR-II from primer 141/901, 116 (77.85%) HGR-III from primer 475/1055, and 126 (84.56 %) HGR-IV from primer 309/1044 (Table 4). In the 86 sequences amplified with primer 465/1048, 25 mutation regions were found, including 21 six consecutive base deletions and ten single base substitutions. For primer 141/901, 37 of the 118 amplified sequences had single or multiple base substitutions. For primer 475/1055, 73 of the 118 amplified sequences had substitutions at 12 sites and 1 insertion. For primer 309/1044, 35 of the 120 amplified sequences had nucleotide substitution, deletion, and insertion mutations (Supplementary S2).

**Table 3:**
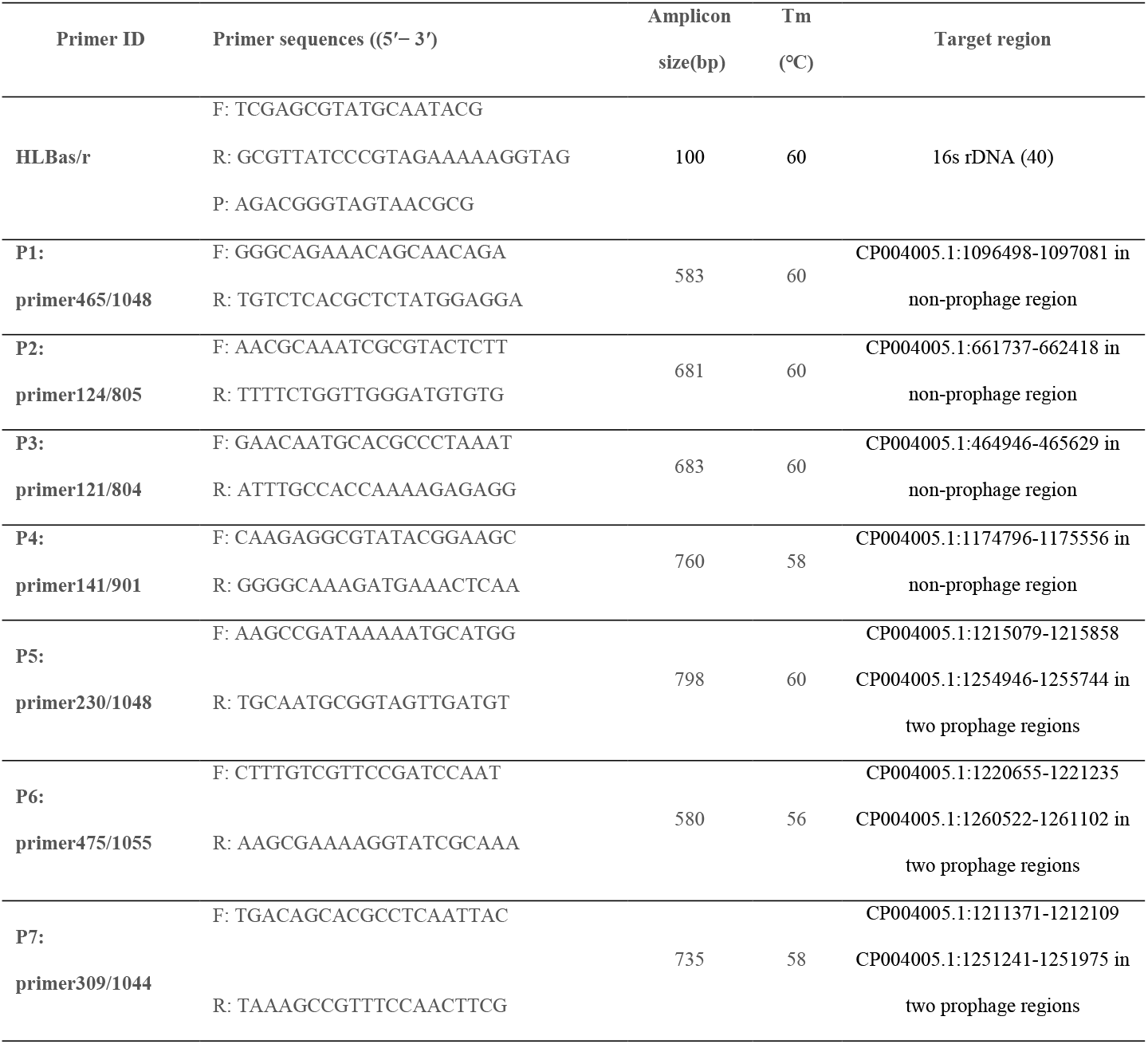
The primer sets used in this study.

**Table 4:** Hypervariable genome regions (HGR) amplified from the *C*Las strains collected in different hosts, symptoms, and geographical locations in Guangxi and Fujian.

|  | Number of<br>samples | Number and percentage (%) of PCR-positive samples |  |  |  | Average<br>amplified (%) |
| --- | --- | --- | --- | --- | --- | --- |
|  |  | HGR-I | HGR-II | HGR-III | HGR-IV |  |
| Geographical origins |  |  |  |  |  |  |
| Yulin, GX | 8 | 5 (62.50) | 8 (100.00) | 5 (62.50) | 5 (62.50) | 71.88 |
| Hezhou, GX | 14 | 12 (85.71) | 13 (92.86) | 12 (85.71) | 12 (85.71) | 87.50 |
| Laibin, GX | 5 | 4 (80.00) | 5 (100.00) | 5 (100.00) | 5 (100.00) | 95.00 |
| Guilin, GX | 16 | 14 (87.50) | 13 (81.25) | 13 (81.25) | 15 (93.75) | 85.94 |
| Wuzhou, GX | 1 | 1 (100.00) | 1 (100.00) | 1 (100.00) | 1 (100.00) | 100 |
| Liuzhou, GX | 12 | 10 (83.33) | 11 (91.67) | 12 (100.00) | 12 (100.00) | 93.75 |
| Nanning, GX | 50 | 40 (80.00) | 39 (78.00) | 34 (68.00) | 42 (84.00) | 77.50 |
| Guigang, GX | 23 | 19 (82.61) | 20 (86.96) | 19 (82.61) | 19 (82.61) | 83.70 |
| Beihai, GX | 10 | 7 (70.00) | 9 (90.00) | 10 (100.00) | 9 (90.00) | 87.50 |
| Sanming, FJ | 10 | 8 (80.00) | 7 (70.00) | 5 (50.00) | 6 (60.00) | 65.00 |
| Hosts |  |  |  |  |  |  |
| Tangerine | 44 | 32 (76.19) | 40 (90.91) | 38 (86.36) | 41 (93.18) | 86.66 |
| Mandarin | 32 | 26 (81.25) | 28 (93.33) | 25 (83.33) | 26 (86.67) | 86.15 |
| Orange | 63 | 55 (90.16) | 48 (76.19) | 46 (73.02) | 52 (82.54) | 80.48 |
| Pomelo | 10 | 7 (70.00) | 10 (100.00) | 7 (70.00) | 7 (70.00) | 77.50 |
| Symptom types |  |  |  |  |  |  |
| Asymptomatic | 44 | 35 (79.55) | 36 (81.82) | 32 (72.73) | 36 (81.82) | 78.98 |
| Yellowing | 11 | 9 (81.81) | 10 (90.91) | 10 (90.91) | 10 (90.91) | 88.64 |
| Blotchy mottle | 30 | 26 (86.67) | 25 (83.33) | 26 (86.67) | 27 (90.00) | 86.67 |
| Zn-deficient<br>like | 64 | 50 (78.13) | 55 (85.94) | 48 (75.00) | 53 (82.81) | 80.47 |
| **Total** | 149 | 120 (80.54) | 126 (84.56) | 116 (77.85) | 126 (84.56) | 81.88 |

All amplicons from 100 HLB positive samples amplified by all four primers were sequenced and aligned. HGR-IV had the most mutation sites, followed by HGR-III, whereas HGR-I was the most conservative region. HGR was least amplified in *C*Las strains collected from Sanming, Fujian, whereas it was highly amplified in those collected from the central parts of Guangxi (Wuzhou, Laibin, and Liuzhou). In the samples from Yulin, Hezhou, and Guigang, the second HGR (HGR-II) was primarily amplified, while HGR-I, HGR-III, and HGR-IV remained stable. In the samples from Laibin, Liuzhou, and Beihai, the amplification rates of HGR-II, HGR-III, and HGR-IV were higher than those of HGR-I. More HGR-I and HGR-IV were amplified from the Guilin and Nanning samples, compared to the HGR-II and HGR-III. HGR-II and HGR-IV were more frequently amplified in HLB strains from tangerine and mandarin, HGR-I from orange, and HGR-II from pomelo. Regarding citrus leaf symptoms, more HGRs were amplified in samples with typical HLB symptoms (blotchy mottle and yellowing) than in samples from asymptomatic and Zn-deficient-like leaves (Table 4).

### Diversity of *Ca*. Liberibacter Asiaticus in China

The SNP loci were detected from four hypervariable genomic regions of 100 samples, including 2190 monophonic loci, 172 polymorphic loci, 69 singleton variable loci, and 103 parsimony loci. A phylogenetic tree constructed from the amplified sequences of four HGR showed that the genetic diversity of *C*Las was not associated with the geographic locations, citrus varieties, and HLB symptoms (Figure 3). One hundred *C*Las strains were divided into four clades. Fifty-seven strains from different geographic sources and citrus species were classified into Clade A (with five reported strains of TaiYZ2, HHCA, HHCA16, YNXP-1, and A4 isolated from citrus) in the United States, Thailand, and China. Thirty-three strains were classified into Clade C with five reported strains of TX2351, GXPSY, CoFLP, JRPAMB1, Psy62 from citrus psyllid and five strains of DUR2TX1, SGCA1, YNJSTC, JXGC, FL17 from citrus. In Guangxi, the remaining ten strains collected from newly expanded citrus groves (Liuzhou, Laibing, Nanning, and Guigang) were clustered into two separate clades without reference strains. GG4 and LZ2 from Mandarin in Clade D had a high variation in the first 170 bp of the prophage regions amplified by primer 475/1055 compared to the other 15 reported genomes. Clade B from Tangerine was quite different from the reference 15 genomes in the prophage regions amplified by primer 309/1044, including LZ7, LZ5, LB1, LZ8, LB3, NN9, GG9, and GG3 (Figure 4).

**Figure 3:**
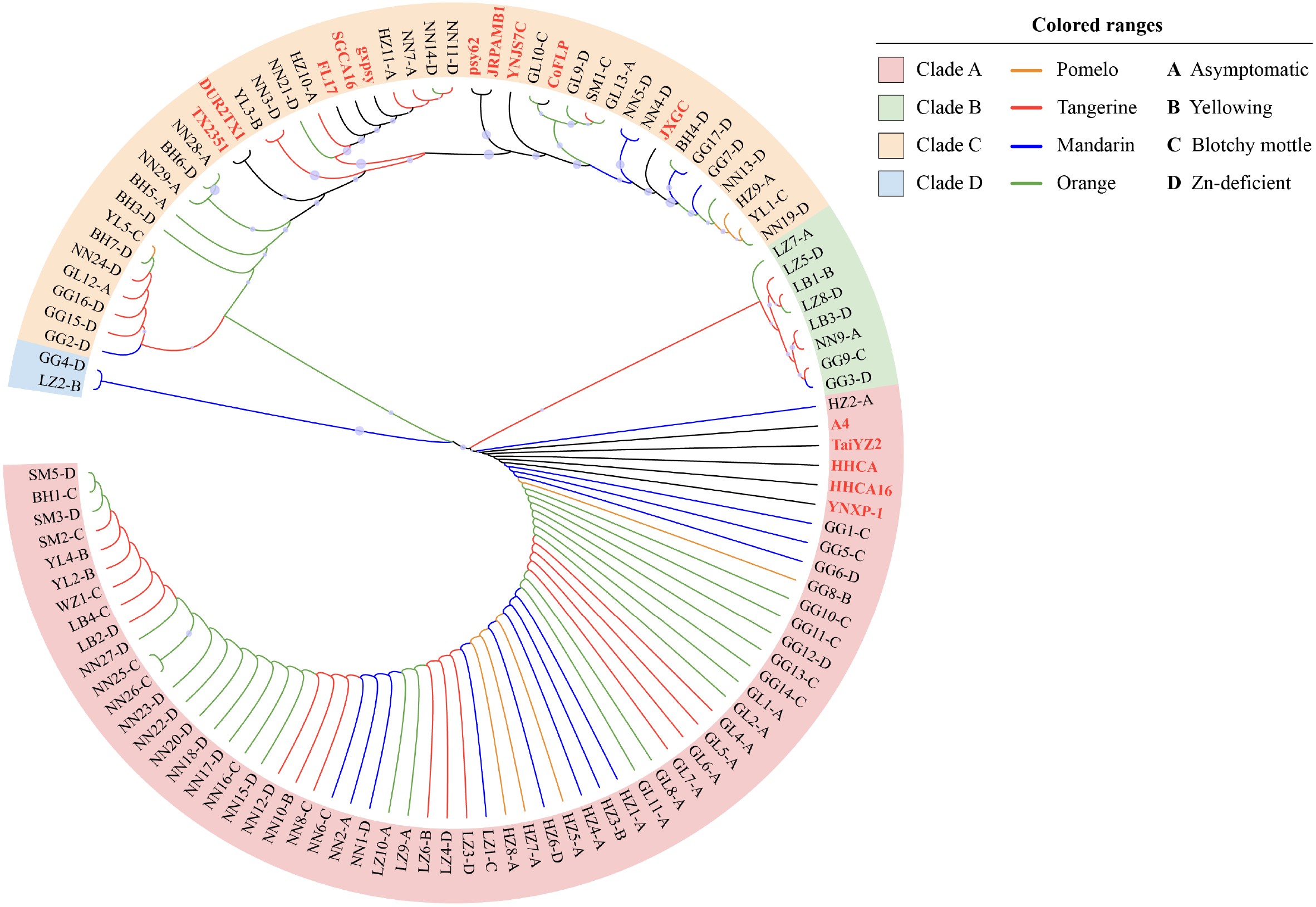
Maximum Likelihood phylogenetic tree based on four hypervariable regions (HGR-I, HGR-II, HGR-III, and HGR-IV) of 100 strains and 15 reported *C*Las. NTSYS clustering was used to analyze the association of *C*Las strains with the citrus geographical origin, varieties, and HLB symptoms.

**Figure 4:**
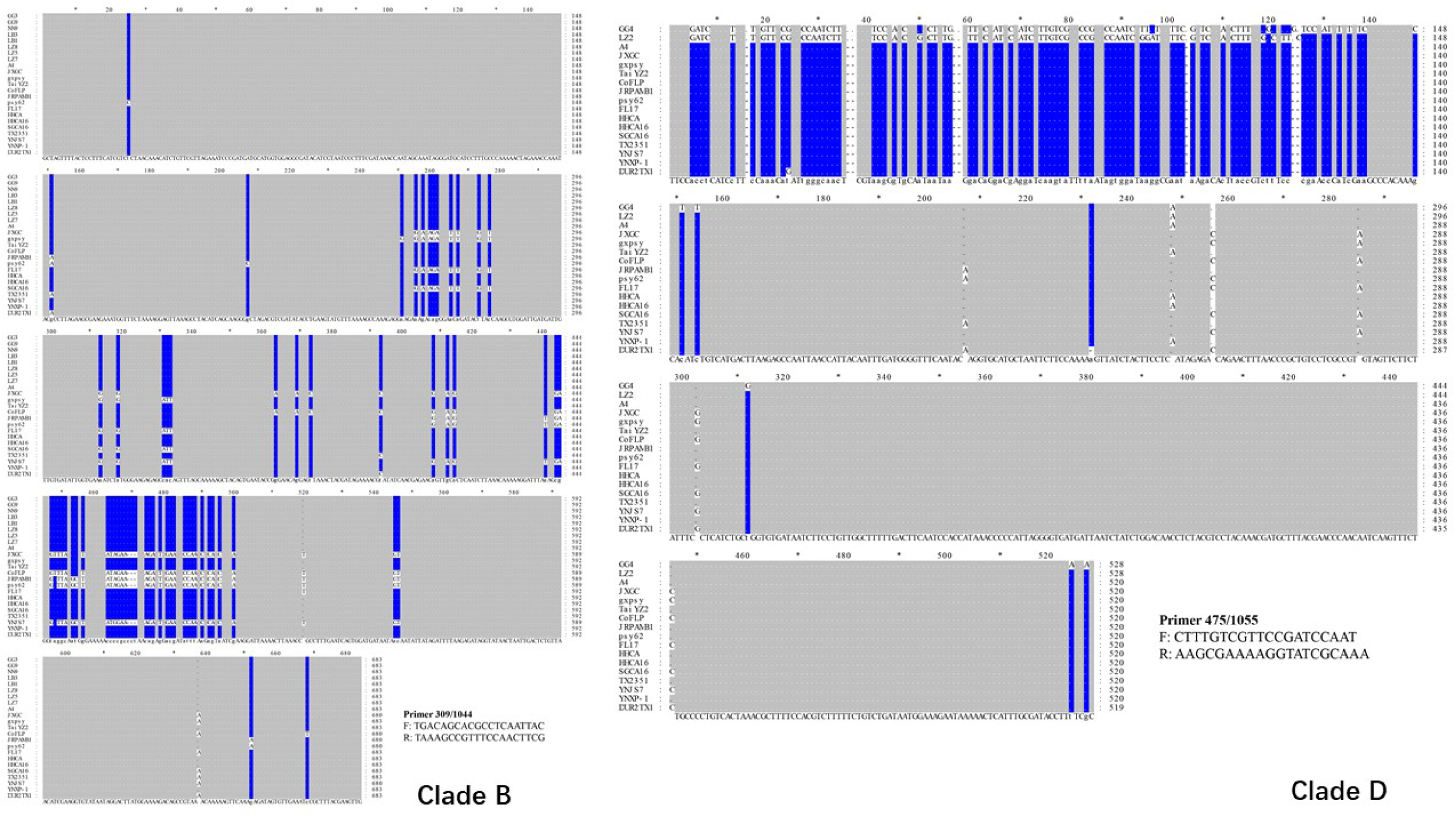
Alignments of the amplified sequences from 10 Clade B and D strains using primer sets 475/1055 and 309/1044 with 15 reported *C*Las genomes downloaded from the NCBI database. The alignments were constructed using DNAMAN. Blue lines indicated the variation sites.

## Discussion

Citrus is the second fruit grown in more than 20 provinces in China, covering an area of more than 2.5 million hectares (16). Citrus production in China has increased by 3.8% annually since 2014, driven by enlarging Guangxi and Sichuan. Guangxi, located at 104° to 112° East Longitude and 20° to 26° North Latitude, has one of the most favorable citrus climates in China, with mild temperatures, adequate light, abundant rainfall, long summers, and short winters (17, 18). Citrus production in Guangxi is now ranked first in China, accounting for 333,000 hectares of planting area and an annual output of 519.3 million tons (19). One of the most serious concerns for the Guangxi citrus industry isCitrus HLB, which is becoming an epidemic and has not been effectively controlled (20). The best way to prevent disease is to eradicate diseased trees and replace them with disease-free seedlings (21).

Citrus HLB is caused by the bacterium ′*Ca*. Liberibacter asiaticus′ (*C*Las) and transmitted by psyllids (22). The incidence of HLB and the distribution of *C*Las were assessed here to determine the potential spread of *C*Las in Guangxi and Fujian, China. A total of 1365 samples were randomly collected from Fujian and Guangxi, with 115 (8.42 %) being PCR-positive. The HLB of the hot and humid climate areas was more severe than that of the cool climate areas. The prevalence of HLB was relatively low in the randomly collected samples from Guilin, (the northern part of Guangxi), where the freezing temperatures in Winter might inhibit the occurrence and spread of the Asian citrus psyllid. The appropriate temperature is conducive to the occurrence and transmission of HLB (23).

Disease symptoms are related to strain type and can be influenced by environmental conditions, internal physiological conditions of the plant, or microorganisms (24). The HLB symptoms in the field were diverse, rendered it difficult to distinguish whether *C*Las caused them or not. HLB-like symptomatic samples had a higher PCR-positive and pathogenic index. Zn-deficient yellowing and blotchy mottle were the most typical HLB symptoms, accounting for more than 40% of the pathogenic index. A detailed examination of some orchards revealed that the orchards on the slope mainly showed blotchy mottle. The orchards with Zn-deficient-like yellowing were observed on the ridge with sparse vegetation.

Citrus varieties are grafted, fused, and heterotypic evolution of citrus gene structure promotes the phenotypic diversity of germplasm symptoms (25). Orange had the highest HLB incidence, with 44.53% of the pathogenic index, followed by mandarin, tangerine, and pomelo. *Poncitus trifoliata* and its hybrids are tolerant to *C*Las (26), exhibit mild symptoms and low bacterial titers, after HLB infection (27). However, grapefruit was found to be susceptible (28–30). Citrus genotypes have been reported to promote variation in *C*Las strains by deciphering *C*Las populations of various citrus varieties (31).

The genetic diversity of *C*Las from different geographical regions and citrus cultivars is vital to predicting the risk of HLB. HLB has been reported to infect all commercial cultivars in China for over one century. However, few genomic variations have been carried out among *C*Las bacterial populations, particularly in Guangxi and Fujian. The results showed that new *C*Las hypervariable genomic region sequences were characterized from 35 previously published genomes. Among the 35 reported genomes, a total of 6,012 single nucleotide variations (SNVs) and 273 small indels were found, with 3,779 variations annotated in protein-coding regions, including 992 synonymous, 2,448 missense substitutions, and 117 frameshift indels. The prophage regions had a higher variation density of 23.3 variations/kb than the non-prophage regions, which had only 3.7 variations/kb. The prophage genes had significantly more synonymous and missense variations than the non-prophage genes. Hypervariable genomic variations (HGR) can lead to the variation of the *C*Las strain population in different hosts and regions (32). Our strains showed significant variations in four HGRs, including two prophages (HGR-III and HGR-IV) and two non-prophage regions (HGR-I and HGR-II). Only 43.44% of the PCR-positive samples amplified one of the HGRs. In the PCR-positive samples from Guangxi and Fujian, the dominant HGRs in each region differed in the amplification frequencies of the HGRs. HGR-II was active in Yulin, Hezhou, and Guigang, while HGR-I was active in Sanming. HGR-II and HGR-IV were more common in mandarin and tangerine isolates, indicating that different citrus varieties have different amplified frequencies. Orange had the higher HGR-I frequency, while pomelo had the highest HGR-II amplification frequency.

Our phylogenetic analysis of four HGRs for *C*Las strains demonstrated that 90% of strains were clustered primarily in two clades, with 15 published genomes from Asia and America, and exhibited genetic diversity. JRPAMB1, Psy62, CoFLp, and Gxpsy isolated from citrus psylla clustered in the same clade as reported in previous studies, while Taiyz2 and A4 isolated from citrus of Thailand and China (in Asia) were also separate. The genotypes of 199 HLB citrus samples collected from Sao Paulo, Parana, and Minas revealed that the genetic variation of *C*Las strains was closely related to geographical location, and environmental factors differed between regions, affecting the genetic diversity of *C*las strains(10). However, *C*Las strains obtained in this study had little relationship with geographical growing regions, citrus varieties, and HLB-like symptoms due to the chaotic nursery stock market in Guangxi, where HLB-scions were planted without quarantine inspection and accelerated *C*Las transmission. Ten strains in two distinct clades considerably varied from previously reported reference *C*Las. Prophages have been found in most *C*Las genomes sequenced so far. However, the origin of prophages from *Liberibacters* associated with plants is not homologous (33). The prophages influence *C*Las pathogenicity, host specificity, and ecological adaptation factors, all of which contribute to the evolution of *C*Las strains. Phylogenetic analysis revealed different *C*Las strains in China, and further research is needed on the biological and phenotypic differences between strains to develop effective disease management and resistance breeding programs.

## Materials and Methods

### Sample Collection

From May 2019 to December 2020, 1789 HLB-like symptomatic and asymptomatic citrus leaf samples were collected from four HLB-affected citrus species in 18 citrus groves in Guangxi and Fujian, China (Figure 5). The leaves of the suspected diseased plants were collected, transported to the laboratory, and stored at - 80°C for DNA extraction. During sampling, geographic coordinates, symptoms, and host citrus species were also recorded to maximize the coverage of *C*Las genetic diversity among citrus samples from various geographical regions and hosts.

**Figure 5:**
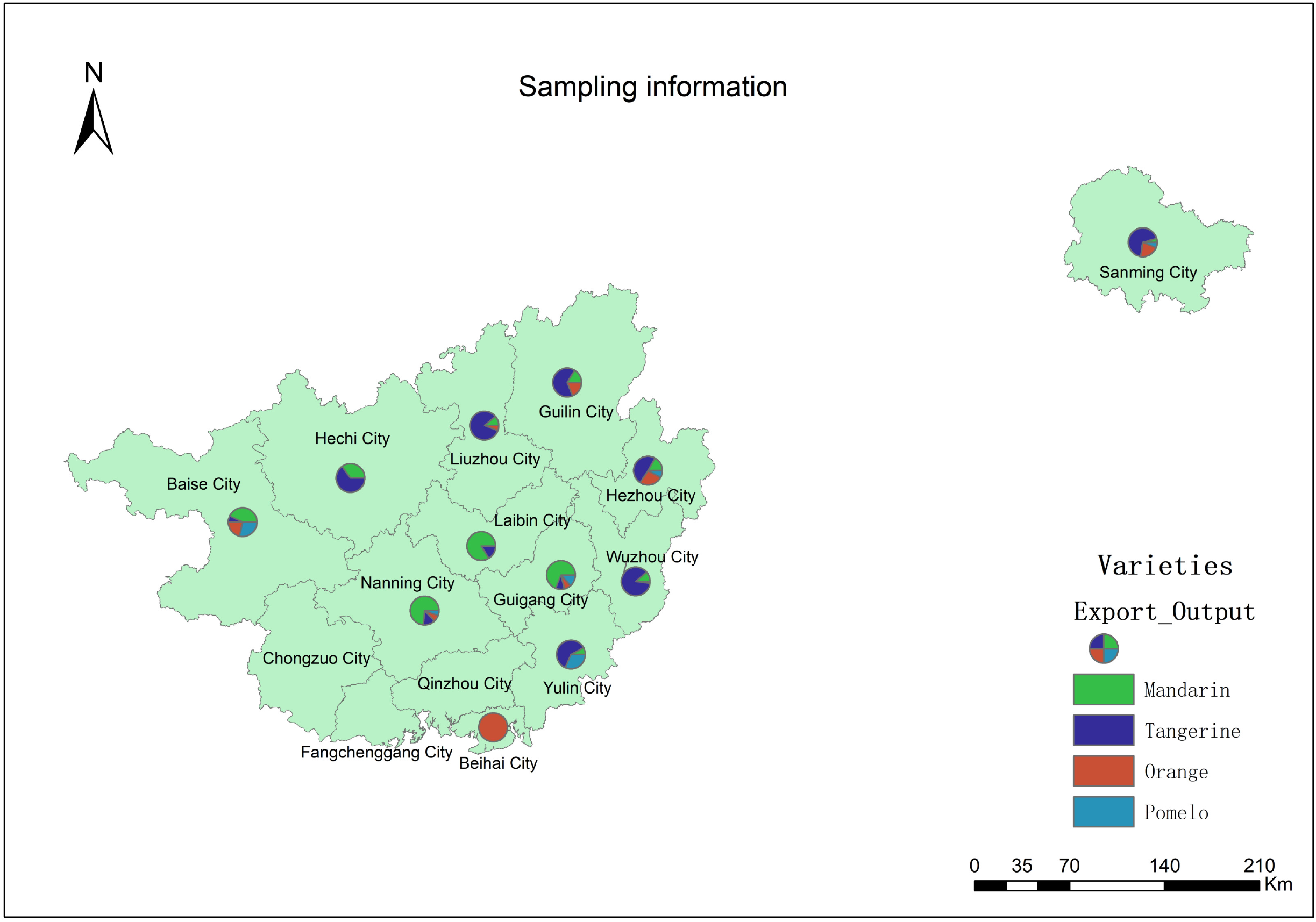
Geographical sites and host plants of the samples collected from Guangxi and Fujian provinces, covering different geographical locations and citrus varieties in China. The map was drawn with Arcgis 10.2 software.

### DNA Extraction

Total genomic DNA from each sample was extracted from the leaf midrib using a modified CTAB extraction protocol. DNA was extracted using 50 μL of TE buffer and stored at −20 °C. The concentration in each sample was normalized to 25 ng/μL after quantifying with a NanoDrop spectrophotometer (IMPLEN, N60).

### Whole-Genome Variation Analysis for *C*Las Bacteria

The genome sequences of GXPSY and 34 other *C*Las isolates were downloaded from the NCBI assembly database with relatively high completeness (>1.0Mb) (Supplementary Table S1). Minimap2 v2.17 was used to align the 34 *C*Las genomes with GXPSY, and the aligned reads were sorted using Samtools v1.12 (34). BCFtools v1.12 called single nucleotide variations (including ≤50bp indels) using the haploid model, which was also used to predict the impact of the variations on gene models (35). BEDTools v2.29 was used to analyze the presence of homologous segments and the density of SNVs and indels across the reference genome in continuous windows (36). Circos v0.69-9 was employed to plot the variation distribution across the reference(37). The nonphage and non-repetitive genomic segments found in all 35 genomes were screened.

Variations with available genotypes were screened in all 35 genomes, and genotypes were concatenated for each before phylogenetic analysis. MEGA v10.2.5 was used to construct a neighbor-joining tree with 1,000 bootstrap replicates using p-distance (38). In Bayesian inference, we ran 1,000,000 generations of Markov chain Monte Carlo (MCMC) tree searches, sampling one tree every 1,000 generations and discarding the first 250 trees. The final phylogenetic tree was a majority-rule consensus tree in which all branches had ≥50% bootstrap support (neighbor-joining) and ≥50% posterior probability (Bayesian inference). Finally, four genomic regions with a high density of SNVs were chosen for PCR amplification and Sanger sequencing. Primer3 v0.4.0 was used to design PCR primers for the different regions shown in Table 1 (39). The primers’ melting temperature (Tm) was set at 60 ± 1°C, and the amplicon length was required to be less than 1kb.

### Primer Design and PCR Assays

qPCR was used to detect the *C*Las bacterium in all leaf samples. Real-time PCR was used to determine the cycle threshold (CT) value employing previously described primer sets and probes (40). Following the *C*Las-infected citrus DNA samples testing, the *C*Las-positive samples were used to amplify seven hypervariable genomic regions using the primer pair (Table 3). The PCR procedure was performed in 25 μL mixtures containing 10 μL of 2× Rapid Taq Master Mix (Vazyme, China), 1 μL forward and reverse primers, 2 μL of template DNA, and 11 μL of H_2_O. The PCR cycles were programmed with an initial denaturation step of 94 °C for 5 minutes, followed by 40 cycles of 95 °C for 30 seconds, 56-60 °C for 30 seconds, and 72 °C for 30 seconds, based on the primer sets used. After the last cycle, a final extension of 72 °C was performed for 10 minutes. Following confirmation, amplification products were stained with ethidium bromide under UV illuminator on 2 % agarose, purified, and sequenced with the amplification primers by Shenggong Biological Engineering (Shanghai Co., Ltd).

### Genotypic and Phylogenetic Analysis

Single nucleotide variations (including ≤50 bp insertion deletions) were invoked in haploid mode using Samtools V1.12 and BCFtools V1.12. Sequence identity and multiple sequence alignments were determined for all DNA sequences using the vector NTI 10 alignment. Single nucleotide polymorphism (SNP) analysis for each sample was carried out using LaunchDnaSP6 software. Specific mutation sites were estimated using Geneious R software (version 9.0.2). Unrooted neighbor-joining (NJ) was used to construct trees, and estimates of chord genetic distances were constructed using MEGA 7. The phylogenetic tree was constructed for the modes of evolutionary divergence, including the *C*Las sequences available in this study and GenBank. The p-distance model was used to calculate genetic diversity differences between populations, and the distance matrix was bootstrapped using 1000 randomizations.

### Statistical Analysis

We calculated the pathogen index (PI) for each location, host cultivar, or HLB-like symptom to reduce background noise in the treated trees. *C*Las bacterial titers were classified into four categories based on CT values, where category 0=CT ≥36.0; category 1=32.0≤CT <36.0; category 2=28.0 ≤ CT <32.0; category 3=24.0 ≤ CT <28.0 and category 4=CT<24.0. For each treatment, the pathogenic index (PI) used to assess the *C*Las bacterial titer was calculated as follows (41).

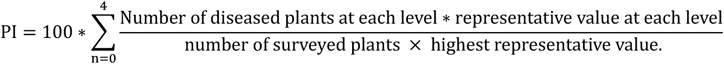

## Acknowledgments

This work was funded by the Key Project of Science and Technology of Guangxi (Guike AA 18118027). We thank all personnel for collecting citrus samples, Jixiu Lin and Jinzhu Zhang (Yongan, Fujian), Xing Gao and Junqi Luo (Nanning, Guangxi), Junyuan Huang (Guangxi University).

## Author Contributions

Muqing Zhang and Chengwu Zou designed research; Fanglan Gao, Wei Yao, and Dean Li collected samples and performed research; Bo Wu and Yixue Bao analyzed data; Fanglan Gao and Muqing Zhang wrote the manuscript; Charles A. Powell and Muqing Zhang edited the paper.

## Competing Interests

The authors declare no conflict of interest with the contents of this article.

## Data availability

All data are included within this manuscript and the Supporting Information. All sequences are deposited in GenBank and listed in Supplementary Table S3.

## Funding and additional information

Guangxi Innovation-driven Development Special Fund Project: Research and Demonstration of comprehensive Control technology of Citrus greening disease (AA18118046)

## Abbreviations

*C*Las: *Candidatus* Liberibacter asiaticus
HLB: Huanglongbing
SNVs: single nucleotide variations
rRNAs: ribosomal RNAs
PI: pathogenic index
CT: cycle threshold
HGR: hypervariable genomic regions
SNP: Single nucleotide polymorphism
CTAB: Cetyltrimethyl Ammonium Bromide
NCBI: National Center for Biotechnology Information
MCMC: Markov chain Monte Carlo
Tm: melting temperature
NJ: neighbor-joining

---

**Supplementary Fig S1:** Gel electrophoresis of the PCR products from P1 primer465/1048 (I), P2 primer124/805 (II), P3 primer121/804 (III), P4 primer141/901 (IV), P5 primer230/1048 (V), P6 primer475/1055 (VI) and P7 primer309/1044 (VII). The amplified products from eight HLB strains were listed in lanes 3–10. Lane 1: Negative control, Lane 2: Positive control, Lane 3:BH6, Lane 4: HZ9, Lane 5:GG7, Lane 6: NN4, Lane 7:GG2, Lane 8:SM10, Lane 9: NN38, and Lane 10: NN24. Lane M L: 2000 Kb ladder.;

**Supplementary Fig S2:** Variation of amplified sequences from hypervariable genomic regions with four pairs of primers, including P1 primer 465/1048 (526bp), P4 primer141/901 (685bp), P6 primer475/1055 (524bp), and P7 primer309/1044 (674bp).

**Supplementary Fig S3:** Minor variation of amplified sequence from non-hypervariable genomic regions with three pairs of primers, including P2 primer124/805 (629bp), P3 primer121/804 (645bp), and P5 primer230/1048 (670bp).

**Supplemental Table S1.** Information of the 35 Las genomes.

**Supplemental Table S2.** The consequence of variations located in genic regions.

**Supplementary Table S3.** The sequence IDs amplified from seven primer sets.

HGR-I, HGR-II, HGR-III, HGR-IV, HGR-V, HGR-VI, and HGR-VII corresponding to different primer sets: primer465/1048, primer124/805, primer121/804, primer141/901, primer230/1048, primer475/1055 and primer309/1044, respectively; “-”: Indicated that the primers did not amplify the sequence.

## Reference

1. Deng X, Lopes S, Wang X, Sun X, Jones D, Irey M, Civerolo E, Chen J. 2014. Characterization of “Candidatus Liberibacter asiaticus” populations by double-locus analyses. Curr Microbiol 69:554–60.

2. Rawat N, Kiran SP, Du D, Gmitter FG, Deng Z. 2015. Comprehensive meta-analysis, co-expression, and miRNA nested network analysis identifies gene candidates in citrus against Huanglongbing disease. BMC Plant Biology 15.

3. Zheng Z, Bao M, Wu F, Chen J, Deng X. 2016. Predominance of Single Prophage Carrying a CRISPR/cas System in “Candidatus Liberibacter asiaticus” Strains in Southern China. PLoS One 11:e0146422.

4. Martini X, Addison T, Fleming B, Jackson I, Pelz-Stelinski K, Stelinski LL. 2013. Occurrence of diaphorina citri (hemiptera: liviidae) in an unexpected ecosystem: The lake kissimmee state park forest, florida. The Florida Entomologist 96:658–660.

5. Sagaram US, DeAngelis KM, Trivedi P, Andersen GL, Lu SE, Wang N. 2009. Bacterial diversity analysis of Huanglongbing pathogen-infected citrus, using PhyloChip arrays and 16S rRNA gene clone library sequencing. Appl Environ Microbiol 75:1566–74.

6. Baldwin E, Plotto A, Manthey J, McCollum G, Bai J, Irey M, Cameron R, Luzio G. 2010. Effect of liberibacter infection (huanglongbing disease) of citrus on orange fruit physiology and fruit/fruit juice quality: chemical and physical analyses. J Agric Food Chem 58:1247–62.

7. Da Graca JV. 1991. Citrus greening disease. Annual review of phytopathology 29:109–136.

8. Yang D, Zeng Z, Zhou L, Li J, Xu C, Zhou L, Wang F. 2019. Identification and Control of HLB Disease in Citrus grandis. Asian Agricultural Research 11:78–82.

9. Islam MS, Glynn JM, Bai Y, Duan YP, Coletta-Filho HD, Kuruba G, Civerolo EL, Lin H. 2012. Multilocus microsatellite analysis of ‘Candidatus Liberibacter asiaticus’ associated with citrus Huanglongbing worldwide. BMC Microbiol 12:39.

10. De Paula LB, Lin H, Stuchi ES, Francisco CS, Safady NG, Coletta-Filho HD. 2019. Genetic diversity of ‘Candidatus Liberibacter asiaticus’ in Brazil analyzed in different geographic regions and citrus varieties. European Journal of Plant Pathology 154:863–872.

11. Davis RI, Gunua TG, Kame MF, Tenakanai D, Ruabete TK. 2005. Spread of citrus huanglongbing (greening disease) following incursion into Papua New Guinea. Australasian Plant Pathology: AAP 34:517–524.

12. Kazan K, Manners JM. 2009. Linking development to defense: auxin in plant-pathogen interactions. Trends Plant Sci 14:373–82.

13. Zuniga C, Peacock B, Liang B, McCollum G, Irigoyen SC, Tec-Campos D, Marotz C, Weng NC, Zepeda A, Vidalakis G, Mandadi KK, Borneman J, Zengler K. 2020. Linking metabolic phenotypes to pathogenic traits among “Candidatus Liberibacter asiaticus” and its hosts. NPJ Syst Biol Appl 6:24.

14. Pitino M, Hoffman MT, Zhou L, Hall DG, Stocks IC, Duan Y. 2014. The phloem-sap feeding mealybug (Ferrisia virgata) carries ‘Candidatus Liberibacter asiaticus’ populations that do not cause disease in host plants. PLoS One 9:e85503.

15. Zhou L, Powell CA, Hoffman MT, Li W, Fan G, Liu B, Lin H, Duan Y. 2011. Diversity and plasticity of the intracellular plant pathogen and insect symbiont “Candidatus Liberibacter asiaticus” as revealed by hypervariable prophage genes with intragenic tandem repeats. Appl Environ Microbiol 77:6663–73.

16. Chen J, Zhu Z, Fu Y, Cheng J, Xie J, Yang L. 2021. Identification of Lasiodiplodia pseudotheobromae Causing Fruit Rot of Citrus in China. Plants 10:202.

17. Liu H, Zhang M, Lin Z. 2017. Relative importance of climate changes at different time scales on net primary productivity—a case study of the Karst area of northwest Guangxi, China. Environmental Monitoring and Assessment 189:1–11.

18. Ziqin H, Dongzhang L, Xianda B. 2017. Analysis on the Impact of Mountain Climate on Tourism Development of Ziyuan County, Guangxi. Meteorological and Environmental Research 8:18–21.

19. Huang Y, Chen G, Huang Y, Xiong L, Liu B. 2019. Status of Soil Nutrients in Citrus Orchards of Guangxi. Agricultural Biotechnology 8:150–151, 155.

20. Yanjun G, Tiecheng C, Yibo H, Liying G, Qianhua J, Hui J, Aifang Z. 2021. Prevention and Control Measures of Citrus Huanglongbing and Application Ideas of Repellent Plants. Plant Diseases and Pests 12:17–22.

21. Yang Q, Lan M, Yang H. 2020. The Present Situation of Occurrence, Control and Prevention of Citrus Huanglongbing in Meizhou City, South China. Agricultural Biotechnology 9:30–34, 37.

22. Xiong Y, Liu XQ, Xiao PA, Tang GH, Liu SH, Lou BH, Wang JJ, Jiang HB. 2019. Comparative transcriptome analysis reveals differentially expressed genes in the Asian citrus psyllid (Diaphorina citri) upon heat shock. Comparative Biochemistry Physiology Part D: Genomics Proteomics 30:256–261.

23. Da Graça J, Korsten L. 2004. Citrus huanglongbing: Review, present status and future strategies. Diseases of fruits vegetables volume I:229–245.

24. Passera A, Alizadeh H, Azadvar M, Quaglino F, Alizadeh A, Casati P, Bianco PA. 2018. Studies of Microbiota Dynamics Reveals Association of “Candidatus Liberibacter Asiaticus” Infection with Citrus (Citrus sinensis) Decline in South of Iran. Int J Mol Sci 19.

25. Oueslati A, Salhi-Hannachi A, Luro F, Vignes H, Mournet P, Ollitrault P. 2017. Genotyping by sequencing reveals the interspecific C. maxima / C. reticulata admixture along the genomes of modern citrus varieties of mandarins, tangors, tangelos, orangelos and grapefruits. PLoS One 12:e0185618.

26. Albrecht U, Tripathi I, Bowman KD. 2019. Rootstock influences the metabolic response to Candidatus Liberibacter asiaticus in grafted sweet orange trees. Trees 34:405–431.

27. Boava LP, Sagawa CH, Cristofani-Yaly M, Machado MA. 2015. Incidence of ‘Candidatus Liberibacter asiaticus’-Infected Plants Among Citrandarins as Rootstock and Scion Under Field Conditions. Phytopathology 105:518–24.

28. Fan J, Chen C, Yu Q, Khalaf A, Achor DS, Brlansky RH, Moore GA, Li ZG, Gmitter FG, Jr. 2012. Comparative transcriptional and anatomical analyses of tolerant rough lemon and susceptible sweet orange in response to ‘Candidatus Liberibacter asiaticus’ infection. Mol Plant Microbe Interact 25:1396–407.

29. Folimonova SY, Robertson CJ, Garnsey SM, Gowda S, Dawson WOJP. 2009. Examination of the responses of different genotypes of citrus to huanglongbing (citrus greening) under different conditions. Phytopathology 99:1346–1354.

30. Halbert SE, Manjunath KL. 2004. Asian Citrus Psyllids (Sternorrhyncha: Psyllidae) and Greening Disease of Citrus: A Literature Review and Assessment of Risk in Florida. Florida Entomologist 87:330–353.

31. Ajene IJ, Khamis FM, van Asch B, Pietersen G, Seid N, Rwomushana I, Ombura FLO, Momanyi G, Finyange P, Rasowo BA, Tanga CM, Mohammed S, Ekesi S. 2020. Distribution of Candidatus Liberibacter species in Eastern Africa, and the First Report of Candidatus Liberibacter asiaticus in Kenya. Sci Rep 10:3919.

32. Hoffman MT, Doud MS, Williams L, Zhang MQ, Ding F, Stover E, Hall D, Zhang S, Jones L, Gooch M. 2013. Heat treatment eliminates ‘Candidatus Liberibacter asiaticus’ from infected citrus trees under controlled conditions. Phytopathology 103:15–22.

33. Gilkes JM, Frampton RA, Smith GR, Dobson RC. 2018. Potential pathogenicity determinants in the genome of ‘Candidatus Liberibacter solanacearum’, the causal agent of zebra chip disease of potato. Australasian Plant Pathology 47:119–134.

34. Li H, Handsaker B, Wysoker A, Fennell T, Ruan J, Homer N, Marth G, Abecasis G, Durbin RJB. 2009. The sequence alignment/map format and SAMtools. Bioinformatics 25:2078–2079.

35. Danecek P, McCarthy SAJB. 2017. BCFtools/csq: haplotype-aware variant consequences. Bioinformatics 33:2037–2039.

36. Quinlan AR, Hall IM. 2010. BEDTools: a flexible suite of utilities for comparing genomic features. Bioinformatics 26:841–842.

37. Krzywinski M, Schein J, Birol I, Connors J, Gascoyne R, Horsman D, Jones SJ, Marra MA. 2009. Circos: an information aesthetic for comparative genomics. Genome research 19:1639–1645.

38. Kumar S, Tamura K, Nei M. 1994. MEGA: molecular evolutionary genetics analysis software for microcomputers. Bioinformatics 10:189–191.

39. Untergasser A, Cutcutache I, Koressaar T, Ye J, Faircloth BC, Remm M, Rozen SG. 2012. Primer3—new capabilities and interfaces. Nucleic acids research 40:e115–e115.

40. Li W, Hartung JS, Levy L. 2006. Quantitative real-time PCR for detection and identification of Candidatus Liberibacter species associated with citrus huanglongbing. Journal of microbiological methods 66:104–115.

41. Yang B, Wang Y, Qian PY. 2016. Sensitivity and correlation of hypervariable regions in 16S rRNA genes in phylogenetic analysis. BMC bioinformatics 17:1–8.

